# Hippocampal neurons construct a map of an abstract value space

**DOI:** 10.1101/2020.12.17.423272

**Authors:** EB Knudsen, JD Wallis

## Abstract

The hippocampus is thought to encode a ‘cognitive map’, a structural organization of knowledge about relationships in the world. Place cells, spatially selective hippocampal neurons that have been extensively studied in rodents, are one component of this map, describing the relative position of environmental features. However, whether this map extends to abstract, cognitive information remains unknown. Using the relative reward value of cues to define continuous ‘paths’ through an abstract value space, we show that single neurons in primate hippocampus encode this space through value place fields, much like a rodent’s place neurons encode paths through physical space. Value place fields remapped when cues changed, but also became increasingly correlated across contexts, allowing maps to become generalized. Our findings help explain the critical contribution of the hippocampus to value-based decision-making, providing a mechanism by which knowledge of relationships in the world can be incorporated into reward predictions for guiding decisions.

## Introduction

Two of the most seminal findings regarding the hippocampus are that hippocampal damage in humans produces a dense amnesia for episodic memory (Eichenbaum, 2013; Scoville and Milner, 1957) and that hippocampal neurons in rodents encode a spatial map of the animal’s environment (Moser et al., 2008; O’Keefe and Dostrovsky, 1971). One way to unify these disparate cross-species findings was the proposal that the hippocampus is responsible for encoding a ‘cognitive map’ (Behrens et al., 2018; Howard et al., 2014; O’Keefe and Nadel, 1978), a network of associations that specifies how various features of the environment relate to one another. Such a map could be used to specify how the various disparate elements of a memory form a discrete episode, or how spatial landmarks form a map of the environment, thereby providing a theoretical framework in which to explain findings in both humans and rodents.

Despite the long-standing claim that hippocampal neurons encode cognitive maps (O’Keefe and Nadel, 1978), the evidence for this assertion is rather weak, relying mainly on the presence of neuronal responses that discriminate non-spatial sensory information (Howard et al., 2014; Ramus and Eichenbaum, 2000; Wood et al., 1999) rather than the representation of a map *per se*. Although a recent study examined the parametric coding of non-spatial information (Aronov et al., 2017), the experimental manipulation was still a sensory stimulus (auditory tone) rather than a cognitive parameter. Recent work in humans, however, have demonstrated hippocampal fMRI signals that are consistent with the encoding of a cognitive map. For example, Theves and colleagues taught humans to categorize abstract stimuli that could be defined in a two-dimensional feature space (Theves et al., 2019). Subsequently, the hippocampal response to the stimuli reflected the relative two-dimensional distances between the objects. In other words, the hippocampus appeared to encode distances in a multidimensional, abstract space the same way that it encoded spatial distances in navigational space.

Although sophisticated paradigms have been developed to explore the cognitive map in humans (Constantinescu et al., 2016; Park et al., 2020; Schapiro et al., 2013; Schuck et al., 2016), neural measures in humans lack the spatiotemporal resolution to determine the precise cellular mechanisms that underlie the representation of the cognitive map. In addition, the behavioral paradigms used in humans are typically too complex to easily translate to the animal model. We recently developed a paradigm that requires animals to track changing reward values associated with sensory stimuli and showed that performance of the task is dependent on the anterior hippocampus (Knudsen and Wallis, 2020), which is the part of the hippocampus that strongly projects to prefrontal areas responsible for processing reward information (Barbas and Blatt, 1995). Reward is potentially a useful variable by which to probe the existence of the cognitive map, since it is an abstract, relational, cognitive parameter that is nevertheless highly salient to animals. We recorded single neuron activity in the anterior hippocampus of two monkeys while they performed this task and found that hippocampal neurons encode precise relationships between reward-predicting stimuli, deemed ‘value place fields’, in the space defined by possible reward values. This supports the idea that the hippocampus does not just encode space, but rather uses a multidimensional code to encode a variety of behaviorally relevant relational information.

## Results

Two macaques (subjects V and T) performed a task requiring them to learn and choose between pairwise combinations of three novel pictures, each associated with the probability of earning a juice reward (Fig. 1A, see Methods) as described previously (Knudsen and Wallis, 2020). Reward contingencies slowly changed throughout the session, requiring subjects to monitor and adapt their choice behavior. The evolving relationship of the pictures’ values relative to one another could be visualized as a trajectory in an abstract 3D value space, with each axis defining the value of one of the pictures (Fig. 1B). Subjects chose the best available picture most of the time (V: 70 ± 2%, 8 sessions; T: 72 ± 2%, 9 sessions) consistent with a simple model-free reinforcement learning (RL) mechanism (mean ± s.e.m. *R*^2^ model vs. behavior; V: 0.48 ± 0.1; T: 0.62 ± 0.06).

**Figure 1.**
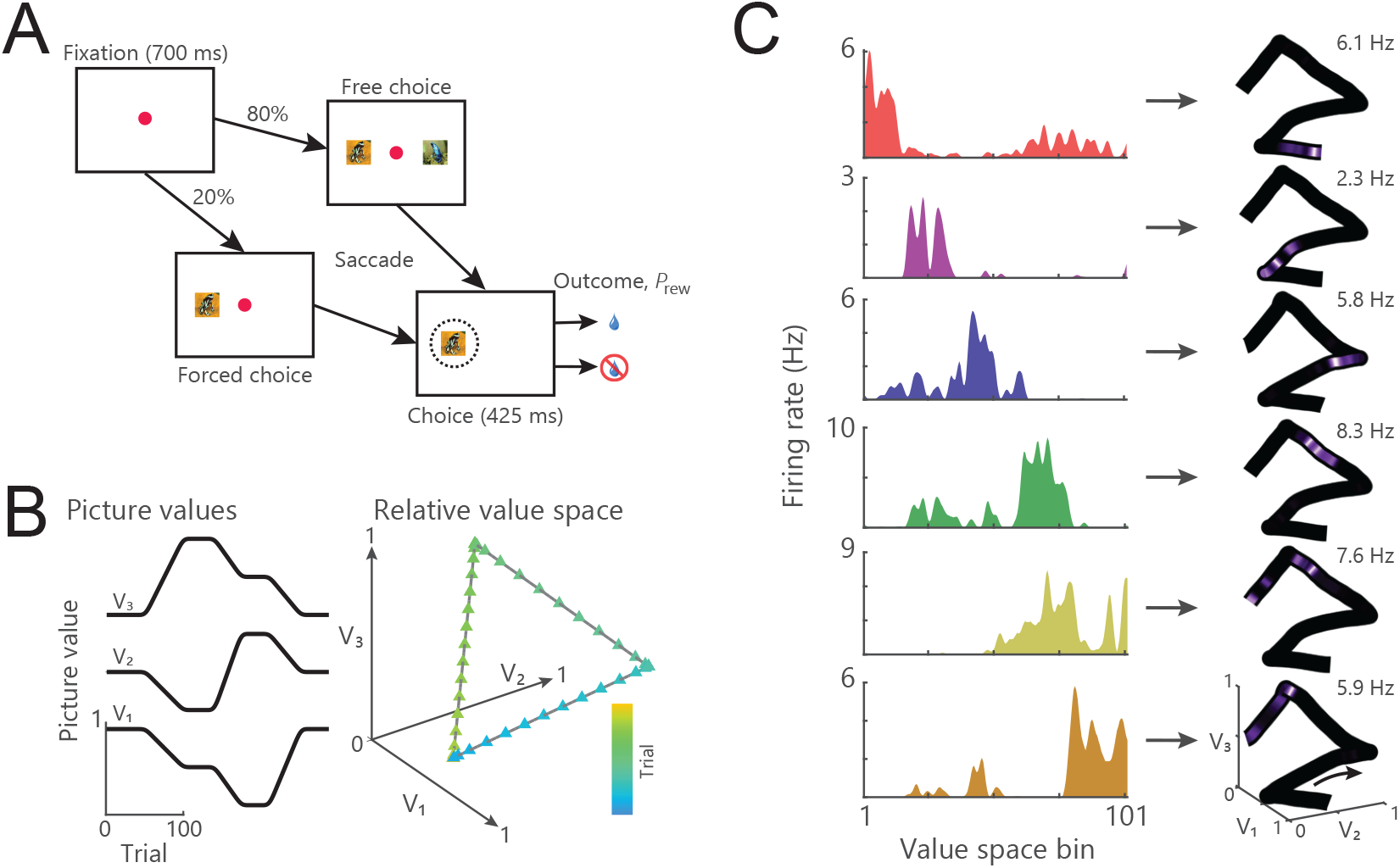
A) Timeline of a single trial. Subjects centrally fixated for 700 ms and were then presented with either a forced or free choice. Subjects selected pictures via saccades, which resulted in the probabilistic delivery of reward (*P_rew_*). B) Example of how the three picture values might change across a session (left) and the resultant trajectory through value space (right). C) Left: Spike density histograms illustrating place encoding in value space for six hippocampal neurons. Right: Firing rates overlaid onto the trajectory through value space mapped from 0 Hz to the peak firing rate as noted in the top right of each panel. Different neurons were active in distinct regions of value space.

We examined the activity of single neurons recorded from the anterior portion of the primate hippocampus (CA3, CA1, Supplementary Fig. 1) during the performance of this task. Many neurons fired bursts of action potentials sporadically over the course of the session. We examined whether this sparse activity encoded task relevant information. We focused on the fixation epoch of the task, since our previous results showed that hippocampal activity during the fixation epoch is critical to task performance (Knudsen and Wallis, 2020). Many neurons appeared to fire at specific places within value space (Fig. 1C). Neuronal encoding during the choice epoch was qualitatively like that in the fixation epoch.

### Hippocampus neurons encode value place fields

To better understand the behavior of these neurons and their relationship to value space, we designed different trajectories to test several predictions. We first examined whether value place fields were replicable by recording from 396 hippocampal neurons (Supplementary Fig. 1; V: 156 neurons, T: 240) as subjects made repeated traversals of a circular path through value space (Fig. 2A). Many neurons appeared to fire at a specific location along the circular path that was consistent across repeated traversals (Fig. 2B). To quantify this, we first identified neurons that had significant value space encoding (see Methods). For each of these neurons (V: 107/156 or 69%; T: 100/240 or 42%), we then correlated value space encoding from the first to the second traversal (Fig. 2C). About 40% of the value selective neurons fired in a consistent region of value space across both traversals (V: 44/107 or 41%; T: 44/100 or 44%; Pearon’s *ρ* assessed at *p* < 0.01). This degree of consistency far exceeded chance levels for shuffled data (Fig. 2C; V: 1.5%; T: 1.3%). Importantly, this consistency in value space was not an artifact of the temporal periodicity of the two trajectories - correlations based solely on time revealed far less consistency between the two trajectories (Fig. 2D; V: 5% significant at *p* < 0.01; T: 3.8%). We analyzed the spatial distribution of fields from one pass to the next (Fig. 2E). There were equivalent numbers of value place fields on both passes (V: 68 vs. 74 fields; T: 76 vs. 78 fields), approximately uniformly distributed along the length of the trajectory through value space. There was no difference in the mean amount of value space information (bits/spike) encoded by the neurons on pass 1 vs. pass 2 (2-sample t-test; V: 0.14 ± 0.02 vs. 0.15 ± 0.01, *t_146_* = −0.18, *p* = 0.86; T: 0.16 ± 0.02 vs. 0.12 ± 0.01; *t_152_* = 1.1, *p* = 0.27).

**Figure 2.**
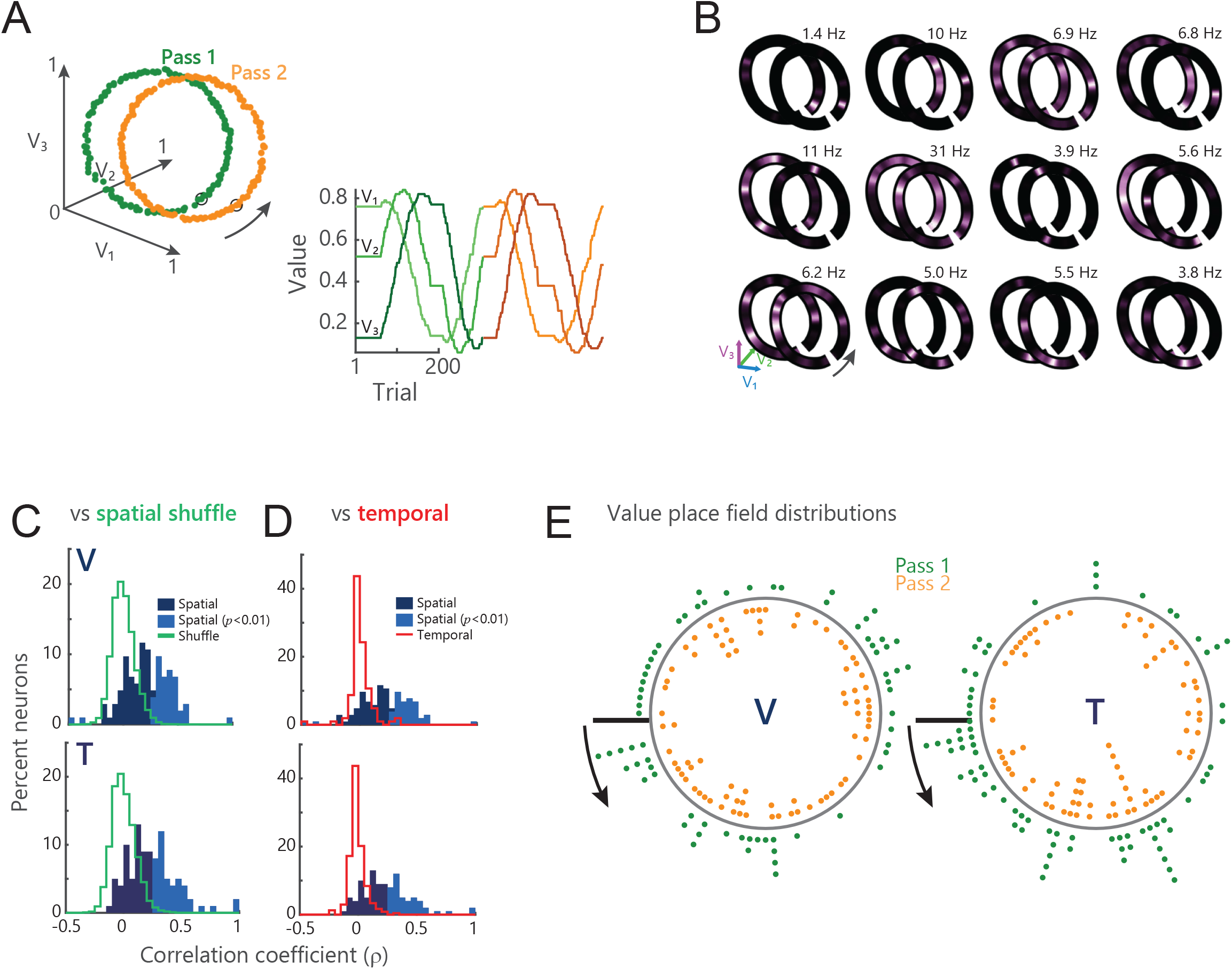
A) Example circular trajectory. Left: picture values plotted in value space. Data points indicate individual trials. Green corresponds to pass 1 and orange to pass 2. Orange trajectory is offset along the *V1* axis for illustrative purposes only. Arrow denotes direction of traversal. Right: picture values plotted in one dimension across the session. Change from green shades to orange shades denote transition from pass 1 to pass 2. Picture values *V1…V3* colored from light to dark shading of given color. B) Examples of hippocampal firing rates mapped on to circular trajectories. Peak firing rates across both traversals are noted in the top right of each panel. C) Distribution of the spatial correlation of firing rates on the first and second traversal for all identified value place neurons. Lighter shades denote correlations significant at *p* < 0.01. Green traces show the distribution of correlations for shuffled data. D) The spatial correlations are the same data as in C), although the y-axis has changed for clarity. The red trace indicates the distribution of temporal correlations for these same value place neurons. E) Distribution of all value place fields on pass 1 (green dots) and 2 (orange dots). Arrows start at value bin 1 (black line) and point in the direction of traversal.

These results suggest that hippocampal neurons are encoding specific locations within value space, and that their value place fields cover the entire value space. However, we also examined whether hippocampal firing might be better explained by other relationships between value parameters, such as the sum of the picture values (Wallis and Rich, 2011). *A priori*, it was unlikely that these factors could account for value place fields, since such tuning would result in clustering of value place fields, rather than the relatively even distribution that we observed. Nevertheless, we explicitly tested for this tuning using a regression model (see Methods, Supplementary Fig. 2). Many neurons did indeed encode these parameters (assessed by *β* values at *p* < 0.01; value sum, value range, value difficulty; V: 32%, 24%, 15% of 156 neurons respectively; T: 36%, 26%, 24% of 240). However, most neurons encoding value place fields did not encode these parameters (V: 73%, 80%, 86% of 107 neurons encoded value place fields but did not encode value sum, value range, or value difference respectively; T 76%, 85%, 83% of 100 neurons).

Value place fields on the circular path were constrained to two dimensions. To test the dimensionality of the value place fields we designed a helical path of similar diameter to the circular path that gradually traversed a third dimension (Fig. 3A). Both subjects completed between 2 and 4 loops of the helix per session, although subject V tended to complete 4 loops more often than subject T (V: 3.2 ± 0.4; T: 2.8 ± 0.3). We assessed value place activity in 578 hippocampal neurons (V: 355, T: 223). About 60% had a place field in at least one loop (224/355 or 63%, in V; 136/223 or 61% in T), with consistent numbers of fields on each loop (chi-squared test, V: loops 1-4, 140/355, 160/355, 115/282, 42/90, *χ*^2^_3,457_ = 2.5, *p* = 0.5; T: loops 1-3, 69/223, 75/223, 78/204, *χ*^2^_3,222_ = 1.2, *p* = 0.5). Many value place fields spanned multiple loops of the helix (Fig. 3B), although the tuning typically became progressively weaker with increased distance from the center of the field (Fig. 3C).

**Figure 3.**
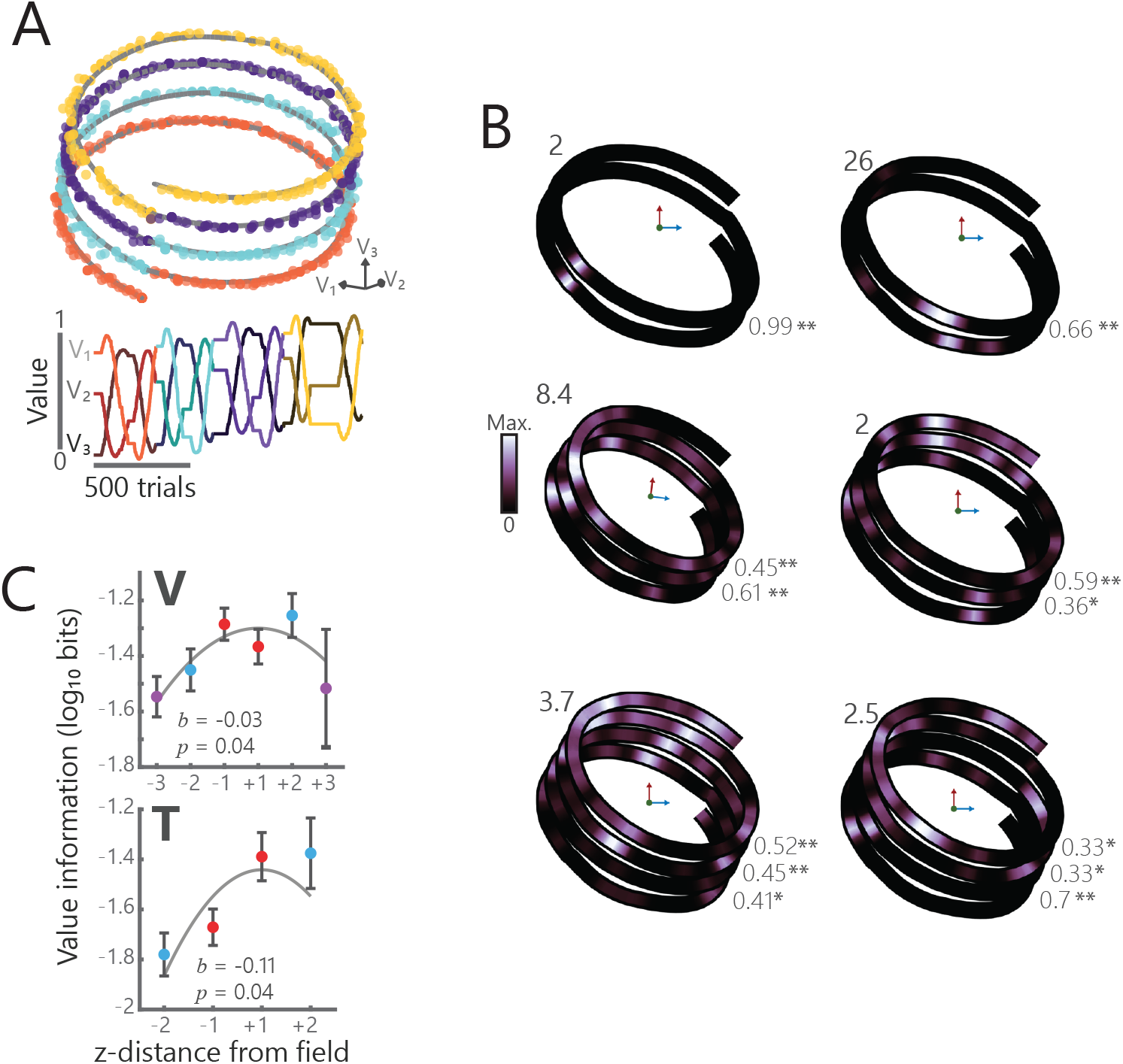
A) Example helical trajectory from a session where subject V performed 4 loops of the helix. Convention follows Fig. 2A. B) Single neuron examples showing value place fields in three dimensions for sessions where subjects completed two (left), three (middle), or four (right) loops of the helix. Numbers in the upper left of each plot denote the maximum firing rate of that neuron. Numbers in the bottom left of each plot denote spatial correlations between adjacent loops for each neuron (* = *p* < 0.01, ** = *p* < 1 x 10^-6^). C) Spatial information of value place fields in three dimensions. For each neuron, 0 was defined as the location with the maximum spatial information and the information content for prior and subsequent loops is illustrated. A quadratic fit of log-transformed spatial information, *I*, by z-distance (*lnI* = *b*_0_ + *b*_1_[z-distance]^2^) was significant in both subjects.

### Value place cells are directionally sensitive

When a rodent alternates runs down a linear track, hippocampal place cells often form directional fields, such that place activity in one direction of travel is largely uncorrelated from the other (Gothard et al., 1996). We tested whether value place fields had similar directional properties by designing a double lemniscate trajectory through value space (Fig. 4A). Subjects made passes through the center of the space from four distal points. This resulted in two passes traversing the central point from the same direction and two passes traversing the central point in opposite directions. Subjects performed the well along this trajectory, choosing the highest value picture most of the time (mean ± s.e.m. optimal choices, V: 68% ± 1%; T: 63% ± 1%)

**Figure 4.**
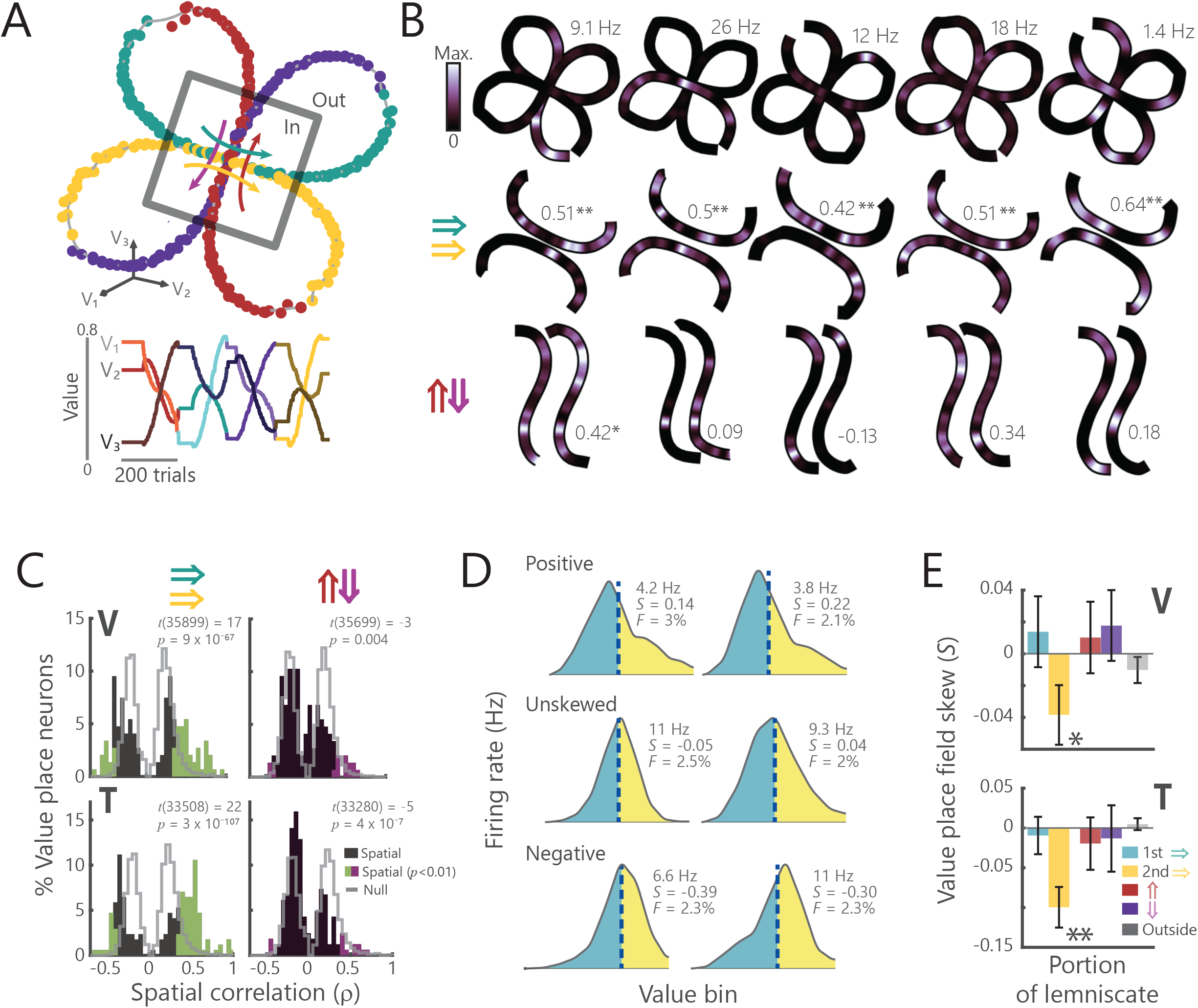
A) Schematic of the double lemniscate task. Convention follows figure 2A. Gray box indicates the overlap regions. B) Five examples of hippocampal neurons encoding position in value space along the double lemniscate trajectory. Top: firing rate for the whole trajectory. The peak firing rate is shown in the upper right. Middle: firing rate on the congruent, rightward passes through the central point. Firing rates are normalized within each pass. Correlation values between the two passes are shown (* = *p* < 0.01, ** = *p* < 0.001). Bottom: firing rate on the opposing (up vs. down) passes. Opposing trajectories have been rotated to be in alignment for visualization. C) Distribution of spatial correlations for congruent (left) and opposing (right) passes through value space for all recorded value place neurons. Colored regions denote significant correlations at *p* < 0.01; gray lines show shuffled null distributions. The proportion of neurons with significant correlations was greater than chance in all cases (two-sample *t*-tests comparing the absolute values of real and shuffled data), but there were significantly more correlated neurons for congruent versus opposing passes (see text). D) Examples of value place fields that were positively skewed (top row; *S* > 0), unskewed (middle row; *S* ≈ 0), and negatively skewed (bottom row; *S* < 0). Vertical dashed lines correspond to the field’s *CoM*. The value *F* denotes the percent of the value space trajectory that the field spanned. Overall, fields typically spanned ~2% of the trajectory (V: 1.8 ± 2%; T: 1.6 ± 1.8%). E) Mean (± s.e.m.) skewness values for different portions of the lemniscate for all value place fields. Significant deviations from zero were determined by one-sample *t-*tests (* = *p* < 0.05, ** = *p* < 0.001. The second traversal of the correlated trajectory resulted in a significant negative skew.

We analyzed 717 hippocampal neurons (V: 365, T: 352) for place activity (see Methods). Consistent with the other trajectories, approximately half of the recorded neurons had a place field on at least one section of the trajectory (Fig. 4B; V: 169 of 365 neurons, 46%; T: 209 of 352, 59%). We correlated place activity along the overlapping portions of the lemniscate, which comprised about 25% of the total trajectory (indicated by the box in Fig. 4A). When subjects passed through the center of space in the same direction, 45% of place neurons were significantly correlated (Fig 4C, V: 55 of 138, 40%; T: 80 of 152, 53%). However, when subjects passed through space in opposite directions, there were significantly fewer correlated neurons (V: 15 of 137, 11%, T: 14 of 149, 9%; chi-squared test; V: *χ*^2^_1,275_ = 30; T: *χ*^2^_1,301_ = 66, both *p* < 0.00001).

Traditional spatial place fields are shaped by experience. They are initially gaussian, but with experience they begin to gradually ramp up in activity as the animal approaches the center of the field, resulting in a negatively skewed place field (Mehta et al., 2000). To examine whether this was the case in our data, for each value place field, we computed a skewness index by comparing firing rates on either side of the field’s center of mass (Fig. 4D, see Methods). Across the population of value place cells, fields were negatively skewed only on the portion of the trajectory that subjects had previously experienced, that is, the second pass along the correlated portion of the trajectory (Fig. 4E). This was consistent with behavioral results which showed that subjects used their prior experience along the trajectory. Subjects’ performance significantly improved on the second pass along the correlated portion of the trajectory relative to the first pass (Supplementary Figure S3).

Taken together, these results show that hippocampal value place cells conserve two important directional properties observed in rodents running on linear tracks. First, neurons are directionally selective. Value space fields are uncorrelated when the subject traverses the same location in abstract value space in different directions but are correlated when the traversal occurs in the same direction. Second, neurons shape their value place fields with experience, as the animal learns to predict the direction of travel along the trajectory.

### Novel pictures remap value place fields

A common feature of hippocampal place fields is that they are not static, but rather can flexibly remap in response to perturbations of the environment. For example, hippocampal neurons may completely shift their preferred firing location in response to changes in one or more features associated with space, or when the location of reward within an environment changes (Anderson and Jeffery, 2003; Fyhn et al., 2007). This global remapping reverses when the original context is restored. We tested if these place cell properties were conserved in abstract value space by introducing a contextual shift in an A-B-A’ block structure using the circular path (Fig. 5A). The two A blocks (A and A’) shared a common set of pictures and the B block used a novel set of pictures while the underlying trajectory through value space was preserved. We hypothesized that the introduction of novel pictures in the B block would require a relearning of stimulus-outcome associations, and hence, would result in a remapping of value place neurons.

**Figure 5.**
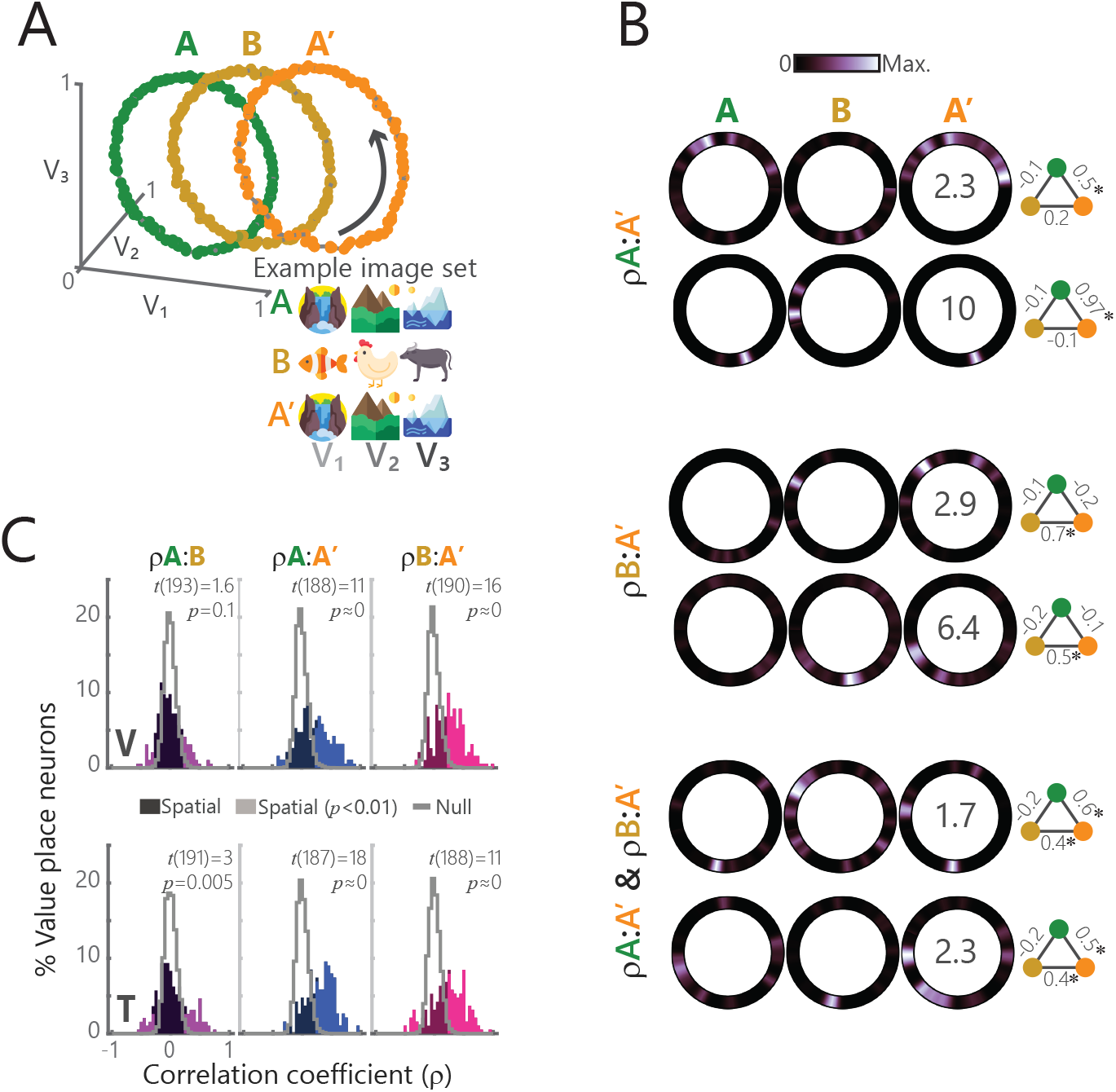
A) Schematic of the ABA’ task. Left: convention follows Fig. 2A. Right: example picture sets for each block. Picture values *V1…V3* follow the trajectories shown in Fig. 2A. B) Single neuron examples showing three distinct patterns of how value place fields change across blocks. Top: two neurons that were significantly spatially correlated on only the A and A’ blocks (*ρ*A:A’). Middle: two neurons spatially correlated on only the B and A’ blocks (*ρ*B:A’). Bottom: two neurons that were correlated on both A:A’ and B:A’ blocks (*ρ*A:A′ & *ρ*B:A′). Numbers superimposed on A’ block activity denote the peak firing rate of the neuron on all three blocks. The schematic to the right of each neuron displays the pairwise correlations (* = *p* < 0.001) between blocks A (green), B (yellow), and A’ (orange). C) Distribution of spatial correlation of firing rates for all recorded value place neurons for all pairwise combinations of blocks. Convention follows Fig. 2C. Test statistics and *p* values from a one-sample *t*-test shown.

We analyzed value place coding in 616 hippocampal neurons (V: 272, T: 344) as described above, quantifying value space fields independently for each block. Many neurons significantly encoded a value place field during at least one block (V: 201 of 272 neurons, 74%; T: 194 of 344, 56%). There tended to be an increased number of identified value place fields as subjects traversed from one block to the next (Cochran’s *Q* test for number of place neurons per block; V: A: 109/272, B: 133/272, A’: 148/272, *Q* = 17, *p* = 0.0002; T: A: 110/344, B: 101/344, A’: 130/344, *Q* = 8.8, *p* = 0.01) suggesting a recruitment of place neurons. For most neurons, the location of the value place field changed from the A block to the B block. We did not find any consistent pattern in this remapping. However, when the task returned to the A block (A’), three distinct patterns emerged. Some neurons switched back to the original field from block A (Fig. 5B, top). Others retained the field from block B (Fig. 5B, middle). Still others retained both fields (Fig. 5B, bottom). The net effect of these changes is that across the population there was little correlation in the location of value place fields between block A and block B, but a strong positive correlation between blocks A and A’ and blocks B and A’ (Fig. 5C).

One possible explanation for these changes in correlational structure across blocks is that they represent the mechanism by which animals can learn to generalize information across blocks to develop a cognitive map of the task. As such, we investigated how experience with the task affected subjects’ behavior and the underlying neural representations. In our previous study, we showed that a model-free RL mechanism (Sutton and Barto, 1998) provided a good fit to subjects’ behavior on the current task when trajectories drifted through the value space at random (Knudsen and Wallis, 2020). In the ABA’ design, where the same trajectories occurred on the B and A’ blocks as the A block, it was possible for the subjects to develop a model of the task that they could use to predict how the reward contingencies would change (Mark et al., 2020). If this were the case, we would expect the fit of the model-free RL mechanism to steadily worsen as the subjects learned the task model.

To examine this, we performed a 2-way ANOVA with the model-free RL fits as the dependent variable and independent variables of Block (A, B, or A’) and Session (Fig. 6A). There were significant Block x Session interactions in both subjects (V: *F_8,1495_* = 523, *p* = 0; T: *F_6,1195_* = 338, *p* = 0). An analysis of the simple effects in subject V showed that model-free RL fits became progressively worse with experience for all three blocks (Block A: *F_4,495_* = 276, *p* = 0.04; Block B: *F_4,495_* = 997, *p* = 0.02; Block A’: *F_4,495_* = 2105, *p* = 0.02). This is consistent with subject V learning how to apply the trajectory during block A to blocks B and A’, thereby worsening the ability of model-free RL to explain the subject’s choice behavior in those blocks. In subject T, the simple effects analysis attributed the Block-Session interaction to a worsening fit with experience on Block A (*F_3,395_* = 1134, *p* = 0.02), but not Blocks B or A’ (B: *F_3,395_* = 43, *p* = 0.11; A’: *F_4,395_* = 22, *p* = 0.15). This may have been because subject T, unlike subject V, received training on the task prior to recording, such that he had already acquired a task model by the time we recorded. Indeed, an analysis of subject T’s training data revealed a pattern like subject V (Supplementary Figure S4). We next examined how these changes in learning strategy mapped onto the neural representation. In subject V, in conjunction with the reduction in model-free learning, there was a significant increase in the strength of correlations of value place neurons between A to A’ and B to A’ and an increase in the number of value place neurons (Fig. 6B). These changes were not evident in subject T, who had been trained on the task structure prior to recording.

**Figure 6.**
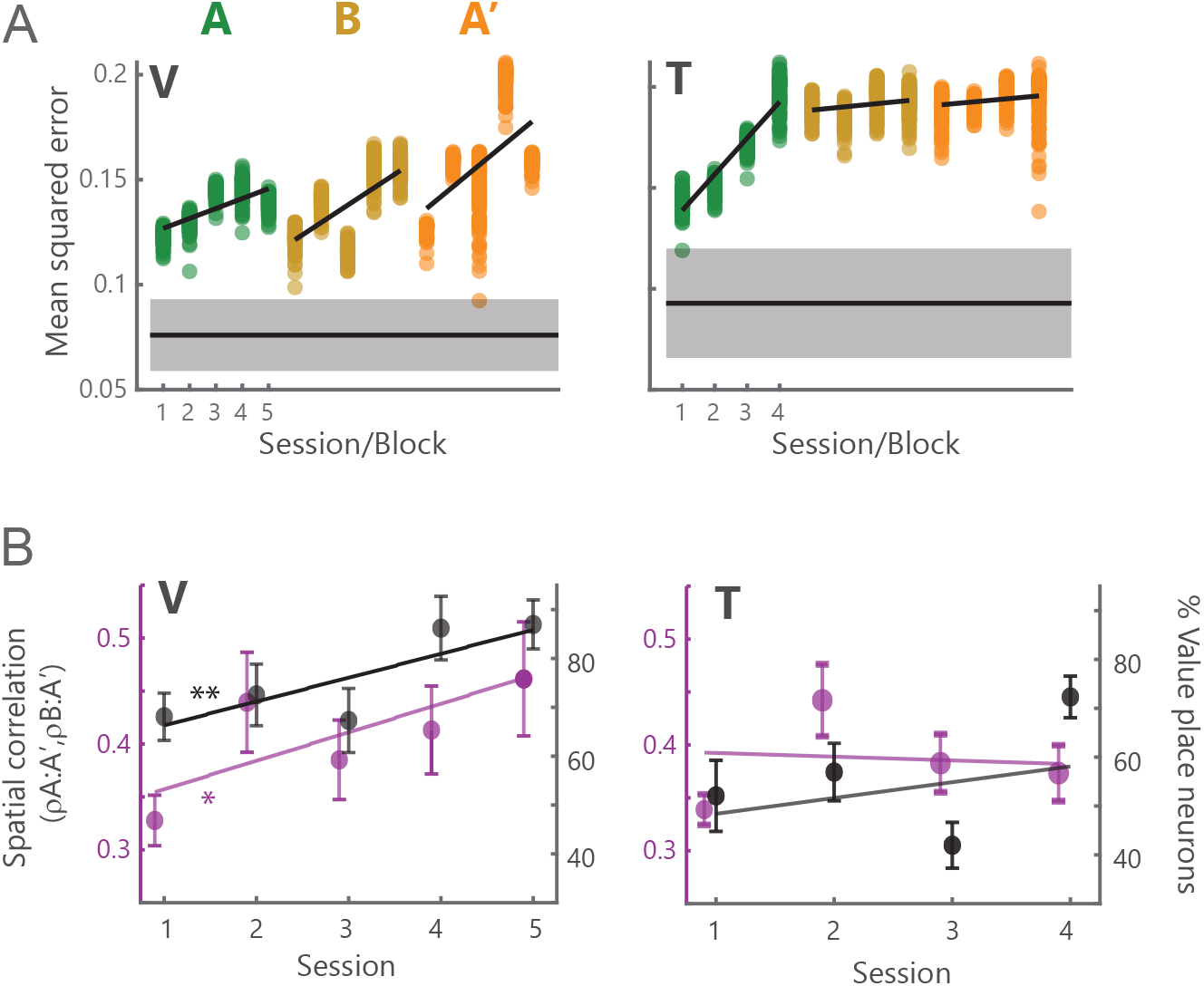
A) Mean squared error between subjects’ choice behavior and model-free RL fits. Black horizontal line and gray shaded regions correspond to the mean (± s.e.m.) for this same analysis when the same subjects traversed arbitrary trajectories through value space (Fig. 1). Each dot (100 per session) corresponds to a single model-free RL simulation of choice to pseudorandomized presentation of stimuli along the circular trajectory using model parameters that were based upon best fits to behavioral data. B) Mean (± s.e.m.) correlation of A:A’ and B:A’ correlated neurons (left axes, purple traces) and overall proportion of value place neurons (right axes, black traces) across sessions (linear regression, * = *p* < 0.05, ** = *p* < 0.01,).

In summary, value place fields were not static but showed flexibility that appeared to correlate with the animal’s experience with the trajectory. Novel pictures induced a remapping that was retained even when the original pictures were restored. These stored maps could explain the ability of the subjects to abandon model-free learning strategies as they generalized across picture sets to learn the underlying trajectories.

## Discussion

A fundamental goal of neuroscience is to understand how the brain constructs internal representations of the world to flexibly guide behavior. In the Euclidean world, hippocampal place cells form one component of such a representation. However, whether the problem at hand is navigating to the coffee shop or navigating your social network when you arrive, cognitive maps, spatial, sensory, and conceptual alike, appear to share a common computational substrate for the systematic organization of structural information (Behrens et al., 2018). Here, we present findings that single neurons in the primate hippocampus construct a map for an abstract cognitive variable through place-like representations. Specifically, the map identifies the relative value of three choice options. Value place neurons exhibited four canonical properties of their Euclidean counterparts: consistency across experiences, multidimensional tuning, directional selectivity, and remapping in novel contexts.

### Hippocampal contributions to reward-based learning

Previous studies have demonstrated that the hippocampus encodes reward information in the context of spatial navigation. For example, Sun and colleagues recently found that when rats received reward on every fourth lap around a track, a subpopulation of place neurons were primarily active on one of four laps only, providing an organizational code to track individual episodes in the pursuit of reward (Sun et al., 2020). Other hippocampal signals have also been found to correlate with reward. Spatial information decoded from individual theta cycles predicted a rat’s distance to a reward (Wikenheiser and Redish, 2015), while hippocampal replays decoded from sharp-wave ripples were biased towards rewarded locations (Pfeiffer and Foster, 2013). Causal manipulations have also implicated a role for the hippocampus in reward processing. Humans with hippocampal damage have difficulty making reward-based decisions (Bakkour et al., 2019), while microstimulation of the anterior hippocampus at the theta frequency disrupts reward-based learning (Knudsen and Wallis, 2020). Our results suggest the specific neural code that might underlie these deficits. Value, like space, is an inherently relational construct (Louie et al., 2011; Rangel and Clithero, 2012): we determine the value of a reward relative to other potential outcomes. For example, the exact same reward can be experienced as either positive or negative, depending on whether an alternative choice would have led to a larger or smaller reward (Camille et al., 2004). Hippocampal activity appears optimized for representing a relational code (Eichenbaum et al., 1999; Whittington et al., 2020), and so it appears that it is used for representing a map of the relative value of options to one another, just as it can also be used for representing a map of the relative location of objects to one another. Indeed, results from neuroimaging in humans have suggested that the hippocampus may be broadly implicated in representing abstract relationships in cognitive space (Park et al., 2020; Theves et al., 2019).

Computational accounts have described two distinct mechanisms by which reward outcomes can be learned (Dolan and Dayan, 2013). Model-free RL is associated with habits and skills, and relies on trial-and-error, storing or caching the values of past actions, and inflexibly repeating those actions that led to higher values. Model-based RL is associated with goal-directed actions and generates predictions via a computationally expensive process that depends on a model of the task at hand but is also able to flexibly respond to environmental changes. There is substantial overlap between the concept of a task model and a cognitive map (Schapiro et al., 2013; Wikenheiser and Schoenbaum, 2016). Two brain regions have been particularly implicated in model-based RL: the hippocampus and the orbitofrontal cortex (Schuck and Niv, 2019; Wikenheiser and Schoenbaum, 2016; Wilson et al., 2014; Zhou et al., 2019). However, it seems unlikely that the orbitofrontal cortex would be responsible for constructing the cognitive map because it typically encodes little information about sensorimotor contingencies (Abe and Lee, 2011; Padoa-Schioppa and Assad, 2006; Wallis and Miller, 2003). This contrasts with the hippocampus where neurons encode sensorimotor contingencies in addition to spatial and temporal contexts, precisely the kind of information that is essential for building task models (Howard et al., 2014; McKenzie et al., 2014). One possibility is that the hippocampus is responsible for constructing the cognitive map that instantiates the neuronal representation of the task model, and orbitofrontal cortex is responsible for using the cognitive map to generate reward predictions that can be used to guide choices. Indeed, we have previously shown that when subjects are performing the same task used in this paper, there is an increase in synchrony in the theta oscillation between orbitofrontal cortex and hippocampus (Knudsen and Wallis, 2020), consistent with a transfer of information between the two structures (Brincat and Miller, 2015). The current results suggest the nature of this transferred information: a concise neural code that represents the value of the potential outcomes relative to one another.

We also considered situations that might influence the geometry of this neural code. First, we found that value place fields represented value space in all three dimensions. In Euclidean space, terrestrial mammals tend to explore their environments in two dimensions, such that place fields are anisotropic even when the rodents move in three dimensions (Grieves et al., 2020), encoding less information in the third dimension relative to the more familiar planar topography. In contrast, place fields in animals who routinely navigate three dimensional space, such as arboreal primates (Ludvig et al., 2004) and aerial bats (Yartsev and Ulanovsky, 2013), display more isotropic fields. Not surprisingly then, a fully abstract space with no cardinal axis, such as value, also yields isotropic fields. Nevertheless, it is possible that manipulations of the value space, such as compressing the variance in one dimension (Conen and Padoa-Schioppa, 2019) may affect how fields are shaped, just as the geometry of a spatial environment determines the shape of two dimensional fields (O’Keefe and Burgess, 1996). We also found that value place neurons are sensitive to the direction of travel through the location in value space. This is consistent with findings in the spatial domain (Gothard et al., 1996; McNaughton et al., 1983) and subsequent findings in non-spatial domains, for example, such as when mice traverse olfactory gradients (Radvansky and Dombeck, 2018). Thus, it is likely that starting from distinct initial states and observing how these states evolve is enough to encourage directional firing. We also found that directionality influences the geometry of the value fields themselves. Opposite direction traversals did not affect field shape, but when a portion of the path was experienced for a second time, fields were negatively skewed. This phenomenon is well documented: as rats make repeated traversals along a linear track, not only do directional place fields emerge with experience (Navratilova et al., 2012), but the fields themselves gradually become negatively skewed (Mehta et al., 2000), reflecting inference about upcoming states.

Although we have drawn parallels between value place fields and spatial place fields, it is likely that the two maps are instantiated by different parts of the hippocampus. In rodents, the dorsoventral axis demonstrates clear functional differences: dorsal neurons have smaller place field sizes and fewer non-spatial representations than ventral neurons (Jung et al., 1994; Royer et al., 2010), while neurons in the intermediate hippocampus fall roughly in between the two poles (Kjelstrup et al., 2008). In contrast, ventral hippocampus is crucial for flexible learning (Avigan et al., 2020), representing task structure (Wikenheiser et al., 2017), and social behavior (Okuyama et al., 2016), high-level behaviors that would benefit from cognitive maps. The dorsoventral axis in the rodent corresponds to the posterior-anterior axis in primates, where there has been a dramatic increase in the size and complexity of the anterior portion of the hippocampus (Insausti 1993), consistent with the sophistication of the primate behavioral repertoire, and the increase in the size and complexity of the frontal cortex, with which the anterior hippocampus connects (Barbas and Blatt, 1995). Thus, we would predict that value place fields may be preferentially located in the anterior hippocampus relative to the posterior hippocampus, whereas the reverse would be true for spatial place fields.

### Generalization of experience

Introduction of novel stimuli while preserving the path through value space induced remapping of value place fields. Similar observations have been made for hippocampal place neurons with respect to physical space (Bostock et al., 1991). For example, remapping of place fields occurred when the color or smell of an arena changed (Anderson and Jeffery, 2003). When the original context was restored, the original place fields returned (Fyhn et al., 2007). Intriguingly, we found that value place field correlations remained high when transitioning from the second context back to the original.

Why might this consolidation of information across contexts occur? Baraduc and colleagues recently observed a similar phenomenon with spatial place fields (Baraduc et al., 2019). Monkeys were trained to forage in two different virtual reality environments, one novel and one familiar. Each environment shared a common underlying spatial structure despite differences in salient landmarks. As the animals learned the novel environment, hippocampal place fields became increasingly correlated with the familiar environment. The authors concluded that this was the underlying neural representation of generalization, whereby a task “schema” is acquired. Consistent with this interpretation, subjects required far less practice in the novel context to achieve good performance, relative to the initial acquisition of the familiar environment. A conceptually similar approach was tested in human participants that learned two image sets corresponding to two different underlying structures (Mark et al., 2020). When participants were given a novel image set in a second session, those who had learned the underlying hidden structure were quickly able to infer relationships between images that were otherwise unknown, suggesting the formation of a schema.

In our task, the subpopulation of neurons that showed consolidated maps across contexts could therefore provide a stimulus-independent description of the structure of the path through value space. Consequently, when the picture set changed, subjects could quickly infer their starting location in value space and use this to predict how the reward contingencies associated with the pictures would change across trials. This could obviate the need to learn the reward contingencies through trial-and-error sampling. Indeed, when we fit behavior to a trial-and-error learning model, our ability to predict behavior decreased as subjects had repeated exposures to the structure of the ABA’ task.

### Future directions

The results presented here demonstrate the first direct neurophysiological basis of a hippocampal map in a purely cognitive space, providing a crucial link between studies of relational memory and spatial navigation. We also showed that the value space map shares many of the features of the hippocampal map of Euclidean space. This raises the question whether other hippocampal mechanisms, such as replay (Foster, 2017; Foster and Wilson, 2006), theta cycling (Kay et al., 2020), and phase precession (Mehta et al., 2002) are also evident in hippocampal value encoding. Evidence of replay-like activity has been identified in the human brain during the performance of nonspatial tasks (Liu et al., 2019; Schuck and Niv, 2019), but it is not known whether this effect is driven by populations of hippocampal neurons. Furthermore, it is unknown whether value place cells interact in the larger hippocampal-entorhinal system. Activity consistent with grid cells has been demonstrated in the entorhinal cortex in a variety of domains, including olfactory information in the rodent (Bao et al., 2019), spatial information in the monkey (Killian et al., 2012), and conceptual information in humans (Constantinescu et al., 2016). Future work should examine the extent to which cognitive and spatial codes relay on shared underlying neural codes (Whittington et al., 2020).

## Acknowledgements

This work was funded by NIMH R01-MH117763 and NIMH R01-MH121448. EBK and JDW designed the experiments and wrote the manuscript. EBK collected and analyzed the data. JDW supervised the project. Thanks to M. Yartsev, Z. Balewski, C. Ford, N. Munet, and L. Meckler for useful feedback on the manuscript. The authors declare no competing interests.

## Methods

### Subjects

Two male rhesus macaques (subjects V and T, respectively) aged 7 and 9 years, and weighing 10 and 13 kg at the time of recording, were used in the current study. Subjects sat head-fixed in a primate chair (Crist Instrument, Hagerstown, MD) and interacted with the task via eye movements digitized with an infrared eye-tracking system (SR Research, Ottawa, Ontario, CN). Subjects each had a large unilateral recording chamber centered over the frontal lobe with access to the temporal lobe. All procedures were carried out as specified in the National Research Council guidelines and approved by the Animal Care and Use Committee and the University of California, Berkeley.

### Task Design

Subjects performed a task in which they were required to choose between pairs of pictures (Knudsen and Wallis, 2020). Each picture had a value to the animal that was defined by the probability that its selection would be rewarded with juice. A single trial started with the presentation of a small, red fixation cue in the center of the screen. Subjects were required to fixate this cue for 700 ms (‘Fixation’ epoch). Following fixation, either one (forced choice trials, 20%) or two (free choice trials, 80%) pictures were presented at 8° of visual angle to the left or right of the fixation cue. Picture locations were pseudorandomized and counterbalanced. Subjects were free to visually inspect each available option and ultimately indicated their choice by maintaining fixation on the chosen picture for 425 ms. Following choice, a delay of 500 ms preceded the delivery of the outcome (reward/no reward). Trials were separated by a 2000 ms intertrial interval. Stimulus presentation and behavioral conditions were controlled using the MonkeyLogic toolbox (Hwang et al., 2019).

The relative values of each of three pictures changed over the course of a session. Furthermore, these changes were designed to map a trajectory through a three-axis ‘value space’ defined by the relationships between the values of the three pictures. For example, the point (0.8, 0.4, 0.3) would correspond to the first picture (*V_1_*) having a value of 0.8 (i.e., rewarded 80% of the time it was selected), *V_2_* a value of 0.4, and *V_3_* a value of 0.3. We designed three trajectories to test various hypotheses: a circle, a helix, and a looping track defined by two orthogonal lemniscates.

To maintain the motivation of the subjects, stable periods, where reward contingencies did not change, were interspersed along the trajectories. These stable points were placed at intervals of *π* for the circular and helical paths, and at the extremities of the lemniscate configuration. Drift periods consisted of the periods between stable points when the reward contingencies changed. We varied the rate of change to ensure that each drift took between 25 and 40 trials. This variability allowed us to decorrelate time and position within value space. The different trajectory configurations were randomly interleaved across recording.

### Neurophysiological recordings

Subjects were fitted with head positioners and imaged in a 3T scanner (Supplementary Figure 1A). The resulting images were then used to generate 3D reconstructions of each subject’s skull and target recording locations within the brain. We implanted customized radiolucent recording chambers over the left hemisphere in both subjects. In each recording session, we lowered up to 4 multisite linear probes into the hippocampus (Supplementary Figure 1B). Electrode trajectories and the appropriate microdrives were defined in software and 3D printed (Form 2, Formlabs, Cambridge, MA) using custom designs (Knudsen et al., 2019). We recorded 2,307 hippocampal neurons (V: 1148, T: 1159) over the course of 27 sessions (V: 16 sessions, T: 11 sessions) using a combination of 16- and 32-channel probes with 100 μm contact spacing (Plexon). Our recordings targeted CA1 and CA3 of the anterior portion of the hippocampus. We confirmed our location in the hippocampus via the presence of sharp wave ripples (Supplementary Figure 1C), prominent theta-band (8-12 Hz) activity (Supplementary Figure 1D), and complex-spiking (bursting) patterns of neuronal activity (Supplementary Figure 1E). Neuronal signals were digitized using a Plexon OmniPlex system with continuous spike-filtered signals (200 Hz - 6 kHz) acquired at 40 kHz and local field-filtered signals (0.1 - 250 Hz) sampled at 1 kHz. We transformed single neuron activity into a binary time series at 1 ms resolution, where 1 indicated the presence of a spike and 0 the absence.

### Behavioral modeling and analysis

To determine how a map of value space might affect choice behavior, we used a standard temporal difference RL model (Sutton and Barto, 1998) that learns picture values *Q_n_* by integrating a prediction error arising from the outcomes of choices:

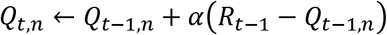

where *Q_n_* is the value of the *n*^th^ picture on trial *t, R* is the outcome of choosing the *n*^th^ picture on trial *t*-1, and *α* is the learning rate, which dictates how much weighting prediction errors have on value updates. Choices are then made by using the standard softmax-activation function:

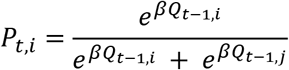

where *Pt,i* is the probability of choosing the picture *i* over picture *j* on trial *t*, and *β* is the inverse temperature parameter that determines the slope of the choice function, and hence the noisiness of choices. Throughout the paper we use the term *V* to indicate the objective value of the picture (the actual probability of reward) and *Q* to indicate the subjective value of the pictures (the animal’s estimate of the value of the picture) derived from the RL model. Optimal fits for both subjects in the TD model were *α* ~ 0.1 and *β* ~ 4, consistent with our previous work (Knudsen and Wallis, 2020).

For the ABA’ task, we fit a model-free RL model to each session (Fig. 6). Using the best fit parameters from subject behavior, we then performed RL simulations of the circular trajectory to assess the accuracy of the model-free account of subject behavior. We subjected the best-fit RL agent (per session) to our experiment by pseudorandomly presenting options along the circular trajectory. For each session, we carried out 100 simulations and quantified error between simulation choices and behavioral choices as mean-squared error.

### Measuring value place fields

Because drift periods were variable in length, our first step was to bin trials to ensure uniformity in our analyses across neurons. We downsampled trajectories to a predetermined number of points; the circular trajectory, individual loops of the helix and passes of the lemniscate path were all tiled at a length of 101 value space bins. Individual trials were then assigned to the appropriate value space bin. Mean neuronal rates during the behavioral epoch of interest were calculated for each bin by dividing the total number of spikes in that bin by the number of trials the subject spent within that bin. For all analyses, these mean rates were used. For visualization, activity was kernel-smoothed with Gaussian kernels (kernel s.d. of 1.2) across bins.

To identify value space encoding, we used information theoretic measures, analogous to those that have typically been used to define spatial place neurons in rats (Skaggs et al., 1993). Thus, the information content of the value space neuron, *I*, in bits per second, is calculated as:

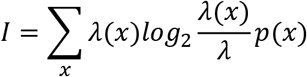

where *x* is the value bin, *λ* is the neuron’s mean firing rate, *λ(x)* is the mean firing rate in value bin *x*, and *p*(*x*) is the probability of being in bin *x* over all trials. We then normalized by *λ* to express *I* in units of bits per spike, a measure of how much value information is conveyed at any location in value space by the neuron. For each neuron, we also calculated a null distribution, by shuffling firing rates 1000 times to preserve overall firing rates but remove correlations with task events. Neurons were considered to have significant value place encoding when three criteria were met: a peak binned firing rate above 1.5 Hz, *I* greater than two standard deviations (95%) above the null distribution, and *I* > 0.2.

We measured the directionality of value place fields in two ways. First, for each field, we determined its center of mass, *CoM*, along the trajectory as:

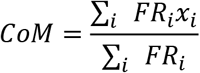

where *FR* denotes the firing rate, *x* is the position, and *i* defines the extent of the field. We determined the skewness of the field, *S*, by generating an index of the mean firing rates to either side of the *CoM*:

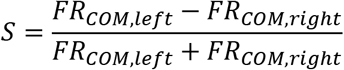

### Calculation of spatial and temporal correlations

To determine the stability of value place fields from one pass through a trajectory to the next, we performed a correlation analysis. For each value place neuron, we correlated the binned firing rates from each pass, using analogous analyses to those that have been used in the rodent literature to study the stability of spatial place fields (Fyhn et al., 2007; Kinsky et al., 2020; Schwindel et al., 2016). For the circular and helical trajectories, we circularly shifted the trajectories relative to one another by ±9 value bins and calculated Pearson’s *ρ* for each shifted pair of passes. We then selected the maximum correlation as the true value. Note that this method can result in strong negative correlations as well as strong positive correlations, due to the circular nature of the data. For example, a neuron might be active in bins 1-5 on the first pass, and 95-100 on the second pass, since these bins are adjacent to one another in value space. To calculate the null distribution of shuffled data, as in Fig. 2C, we shuffled trial-by-trial firing rates and then re-mapped these shuffled rates onto the trajectory through value space, before repeating the correlation analysis. For each neuron and pass, this was repeated 1000 times. We employed a similar shifting procedure for the lemniscate space. However, rather than circularly shifting, we first performed a cross correlation bounded at ±9 value bin lags and computed the Pearson correlation on the lag that produced the largest absolution correlation. This results in fewer correlations at 0 since the procedure is biased towards detecting some correlation. However, the bias is accounted for by our shuffling procedure.

We performed a control analysis to ensure that value space activity was not confounded by a temporal periodicity equivalent to the approximate time taken to complete one pass through the value space. We were able to unconfound this from position within the value space due to variability in the speed at which we changed the reward contingencies during each drift period. To test for temporal periodicity, we correlated the serial order of trial firing rates for each trajectory, truncating whichever pass was longer.

### Testing tuning of hippocampal neurons to other value parameters

We tested hippocampal firing rates for evidence of encoding other relationships related to value, such as those that have been observed in orbitofrontal cortex (Kennerley et al., 2011; Padoa-Schioppa, 2013). Specifically, we defined three additional value-related parameters that could plausibly be encoded: value sum, *Q_sum_*, which is the sum of the value of the three pictures (*Q_1_* + *Q_2_* + *Q_3_*), value range, *Q_rng_*, which is the range of picture values (*Q_max_ - Q_min_*), and value difficulty, *Q_diff_*, which is the discriminability of the two most valuable pictures (*Q_max_* - *Q_mid_*). These regressors were linearly related to mean firing rates during the fixation epoch, *FR_fix_*, as:

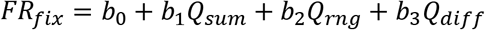

The variance inflation factor for this model was in acceptable bounds for all sessions (<2.5). Significance was assessed at *p* < 0.01.

### Statistics

All statistical tests are described in the main text or the corresponding figure legends. Error bars indicate standard error of the mean (s.e.m.) unless otherwise specified. All comparisons were two-sided.

### Data and Code Availability

The datasets and code supporting the current work are available on request.Supplementary Figure legends

## Supplementary Figure legends

**Supplementary Figure S1.**
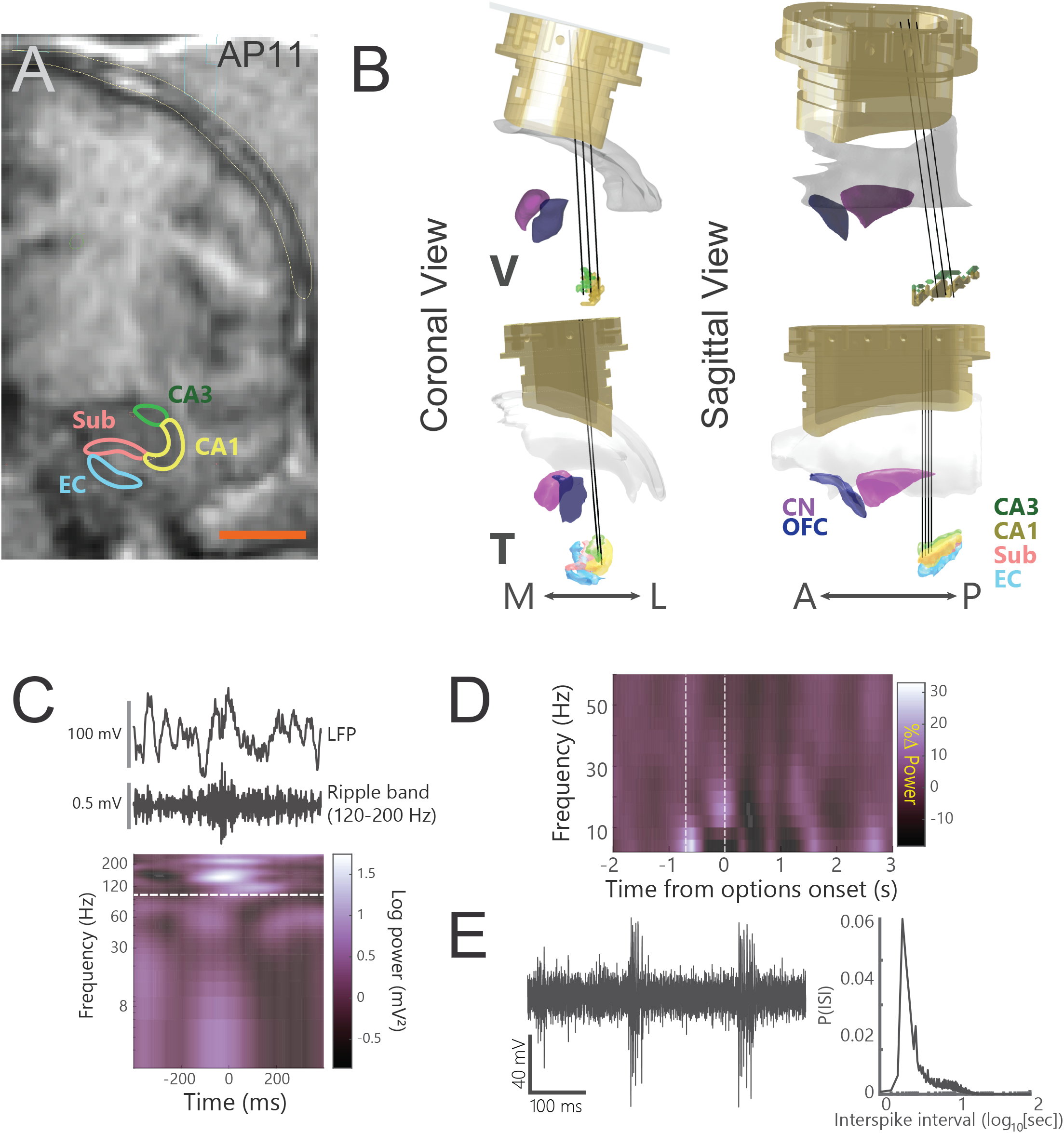
A) Coronal MRI slice from subject T 11 mm anterior to the interaural line, indicating anatomical subdivisions of the hippocampus and surrounding cortex. Subregions defined for modeling are shown. Sub = subiculum, EC = entorhinal cortex. B) Coronal (left) and sagittal (right) views of 3D reconstructions for both subjects. Black lines indicate electrode trajectories into the hippocampus. For each session, probes were offset within a ~200 μm circular window to ensure fresh recording locations. The anterior-posterior extent of the recordings ranged from 11.5 to 14 mm anterior of the interaural line in subject V and 8 to 14 mm in subject T. C) Recordings of the hippocampal local field potential. Top: Raw local field potential from CA1 electrode depicting a sharp-wave event during the intertrial interval. Middle: Frequency-decomposed signal to illustrate activity in the ripple band (120-200 Hz) from the same electrode. Bottom: Normalized local field potential power from 2-250 Hz calculated in 3 Hz steps during the sharp-wave event. White dashed line indicates the low end of the ripple band. D) Hippocampal local field potential power for an example electrode, showing the increase in theta power during fixation. Power is normalized to the mean power during the intertrial interval. E) Left: Raw spike-filtered (1-5.6 kHz) activity for the same electrode in D) showing bursting activity. Right: Interspike interval histogram for the same neuron.

**Supplementary Figure S2.**
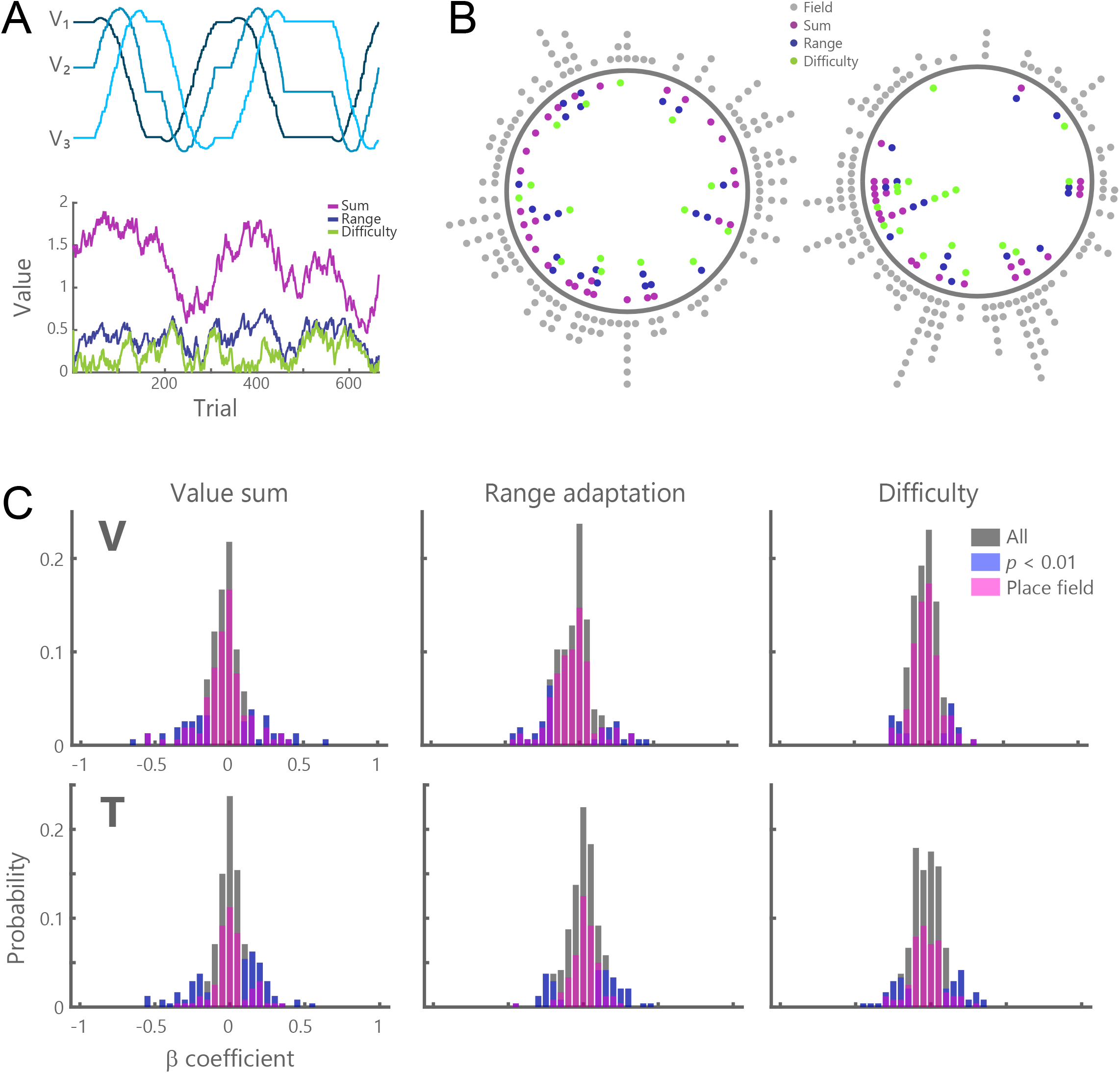
A) Illustration of how value parameters varied from an example session with subject V. Top: individual picture values, as in Fig. 2A. Bottom: parameters of value sum (*Q_1_ + Q_2_ + Q_3_*, purple trace), value range (*Q_Hi_ - Q_Low_*; blue), and value difficulty (*Q_Hi_ - Q_Mid_*, green) derived from subjective value estimates from the same session. B) Relationship between value place fields and other types of value encoding. Colored dots indicate value place fields that also encoded one of the value parameters. Grey dots indicate value place fields that did not encode other value parameters. C) Histograms showing the distribution of regression coefficients for each value parameter for all neurons (gray bars), neurons significantly tuned to a given value parameter at *p* < 0.01 (blue bars), and for all identified value place fields (pink bars). Most value place neurons did not encode other value parameters.

**Supplementary Figure S3.**
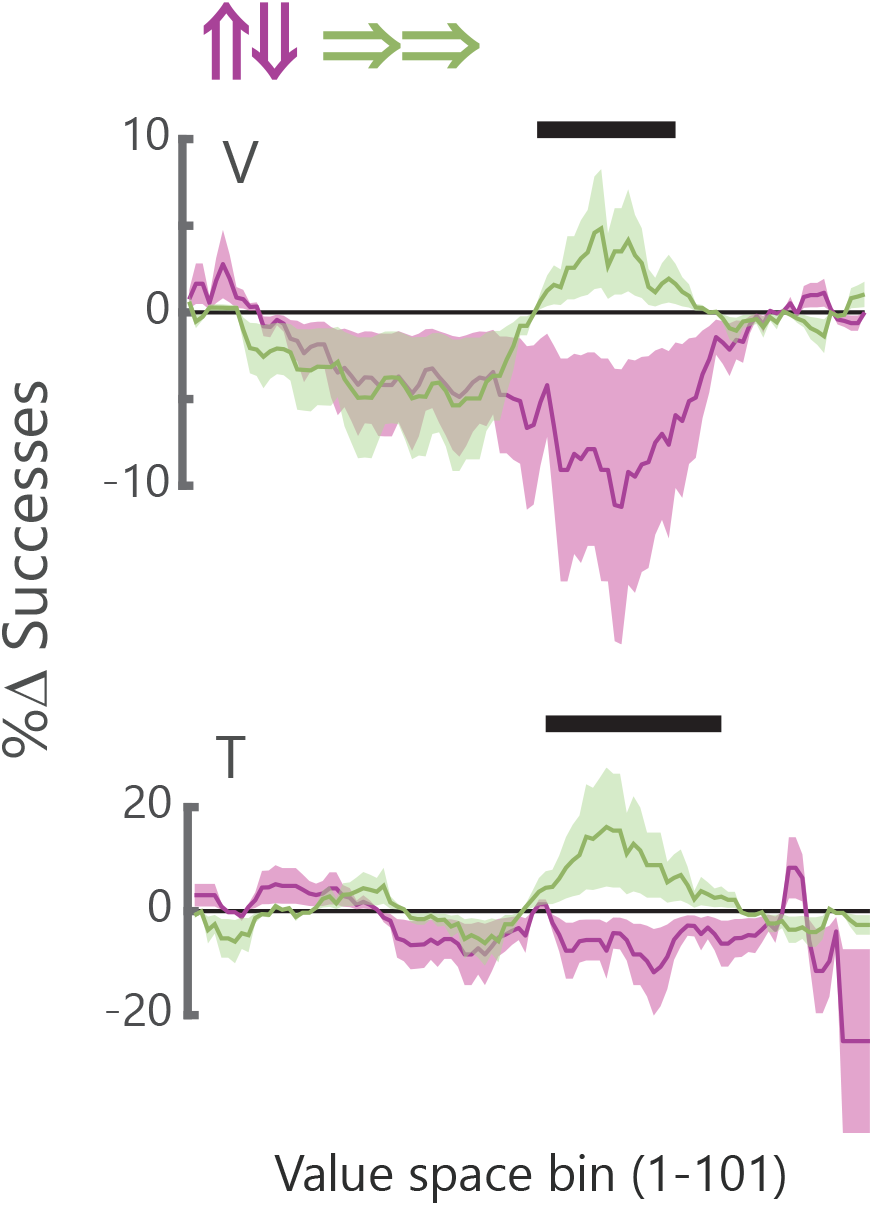
Difference in the mean (± s.e.m.) success rate (probability of choosing the optimal picture) on the second pass through the center location relative to the first pass. Magenta traces are congruent passes (second rightward pass - first rightward pass), while the orange traces are opposing passes (down pass - up pass). Black horizontal lines denote significance at *p* < 0.05 of a sliding paired *t*-test with a window of 6 value bins stepped one at a time. Significance only shown for at least 5 consecutive steps of *p* < 0.05. Performance improves on the second pass relative to the first for the congruent passes, but not the opposing passes.

**Supplementary Figure S4.**
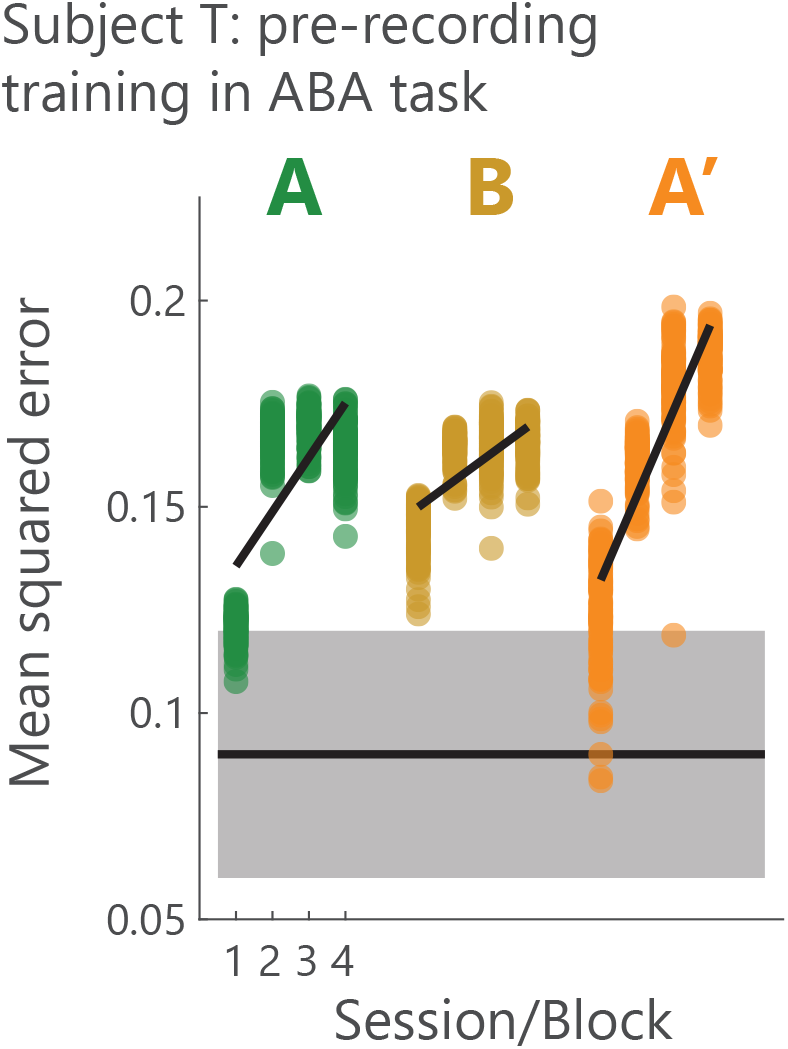
Modeling the pre-recording sessions for subject T when he first learned the ABA’ task structure. Convention follows Fig. 6A. The pattern of results is like subject V when he learned the ABA’ task structure during recording. There was a significant Block x Session interaction (*F_6,1188_* = 233, *p* = 0) and an analysis of the simple effects showed that fits became progressively worse with experience for all three blocks (Block A: *F_3,395_* = 1140, *p* = 0.02; Block B: *F_3,395_* = 230, *p* = 0.05; Block A′: *F_3,395_* = 1654, *p* = 0.02). Thus, both of our subjects appeared to acquire a schema-like representation of the ABA’ task structure at similar rates when they were first exposed to this experimental design, which for subject T was prior to recording and for subject V was during recording.

